# Computational modeling of anthocyanin pathway evolution: Biases, hotspots, and trade-offs

**DOI:** 10.1101/511089

**Authors:** Lucas C. Wheeler, Stacey D. Smith

## Abstract

Alteration of metabolic pathways is a common mechanism underlying the evolution of new phenotypes. Flower color is a striking example of the importance of metabolic evolution in a complex phenotype, wherein shifts in the activity of the underlying pathway lead to a wide range of pigments. Although experimental work has identified common classes of mutations responsible for transitions among colors, we lack a unifying model that relates pathway function and activity to the evolution of distinct pigment phenotypes. One challenge in creating such a model is the branching structure of pigment pathways, which may lead to evolutionary trade-offs due to competition for shared substrates. In order to predict the effects of shifts in enzyme function and activity on pigment production, we created a simple kinetic model of a major plant pigmentation pathway: the anthocyanin pathway. This model describes the production of the three classes of blue, purple and red anthocyanin pigments, and accordingly, includes multiple branches and substrate competition. We first studied the general behavior of this model using a naïve set of parameters. We then stochastically evolved the pathway toward a defined optimum and analyzed the patterns of fixed mutations. This approach allowed us to quantify the probability density of trajectories through pathway state space and identify the types and number of changes. Finally, we examined whether our simulated results qualitatively align with experimental observations, i.e., the predominance of mutations which change color by altering the function of branching genes in the pathway. These analyses provide a theoretical framework that can be used to predict the consequences of new mutations in terms of both pigment phenotypes and pleiotropic effects.

## Introduction

Many complex phenotypes evolve through changes to the activity of metabolic pathways (Morrison and Badyaev 2016; Nam et al. 2011; Pichersky and Gang 2000; Schuster et al. 2000; Strohman 2002). In addition to forming the basis for the extraction and transfer of energy within organisms, metabolism contributes to phenotypes by producing essential cellular products such as structural components, toxins, and pigments (Chen et al. 2014; Colón et al. 2010; Flowers et al. 2007; Keller et al. 2005; Morrison and Badyaev 2017). Evolution of metabolic pathways is known to be shaped by both the topological structure of pathways and the biochemical constraints of individual enzymes (Dykhuizen et al. 1987; Morrison and Badyaev 2017; Nam et al. 2011). Furthermore, the regulatory architecture governing pathway gene expression plays a vital role in the evolution of pathway activity and the resulting alterations in phenotype (Braakman et al. 2017; Gompel et al. 2005; Morrison and Badyaev 2017; Panettieri et al. 2018; Reed and Serfas 2004; True et al. 1999; Wittkopp et al. 2003; Zhang et al. 2017). These properties make metabolic pathways excellent systems to study the fundamental principles that underlie the genotype-phenotype map for complex phenotypes.

A large body of theoretical work has been devoted to understanding the control of metabolic pathways. Metabolic control analysis (MCA) arose from classical enzyme kinetics and solved the problem of analyzing entire metabolic systems simultaneously, rather than focusing on individual components in isolation (Cornish-Bowden 1995; Heinrich and Schuster 1996; Kacser et al. 1995). These methods have been widely used to understand the behavior of metabolic pathways in both basic and applied research (Chen et al. 2014; Colón et al. 2010; Hoefnagel et al. 2002; Schuster et al. 1999, 2000; Stephanopoulos 1999). Still, few studies have used MCA to investigate the evolution of metabolic pathways and the resulting phenotypes, and these have focused on simple pathways with little or no branching (Heckmann et al. 2013; Rausher 2013; Wright and Rausher 2010). While the results point to predictable evolutionary patterns, such as the concentration of fixed mutations in enzymes with the most control over pathway flux, it remains unclear whether these ‘rules’ apply to the large and highly branched metabolic pathways common in biological systems. Such complex branched topologies are likely to generate trade-offs due to competition for substrates and enzymes (Dekel and Alon 2005; Dykhuizen et al. 1987), and thus may introduce constraints on evolutionary trajectories (Guillaume and Otto 2012).

Although theory has not extensively examined the evolution of phenotypes arising from branching pathways, empirical studies can provide insight into the types of mutations that contribute to evolutionary change. Evolutionary studies on experimental systems, such as carotenoid metabolism in birds (Morrison and Badyaev 2016, 2018), various metabolic pathways in yeast (Wisecaver et al. 2014) and Drosophila (Flowers et al. 2007), and anthocyanin biosynthesis in flowering plants (Smith and Rausher 2011; Streisfeld and Rausher 2009a,b), have demonstrated that causal mutations can be drawn from several broad classes, such as regulatory vs. biochemical (structural) and upstream vs. downstream. Depending on the genetic architecture, these classes tend to be differentially favored during evolutionary trajectories, leading to detectable genomic “hotspots” for phenotypic evolution (Cooper et al. 2003; Cresko et al. 2004; Esfeld et al. 2018; Martin and Orgogozo 2013; Treves et al. 1998; Woods et al. 2006). Pleiotropic effects have commonly been suggested as an explanation for these patterns (Gompel et al. 2005; Streisfeld and Rausher 2009b; Wessinger and Rausher 2014) although other factors, such as target size and chromosomal location, may also play a role (Arbeithuber et al. 2015). Determining how pathway structure gives rise to these predictable patterns is critical for understanding the evolution of the phenotypes that result from metabolic activity (Kuang et al. 2018; Wessinger and Rausher 2012b, 2014).

Flower color has proven to be a particularly useful model for studying the origin of novel phenotypes through metabolic evolution. Coloration in plants is primarily determined by the presence of various pigment compounds, the most common of which are produced by the highly conserved anthocyanin pathway (Fig. 1, S1). This pathway is part of the larger flavonoid biosynthesis pathway, which produces an array of related flavonoid compounds, that includes the anthocyanins and flavonols. These compounds play roles as pigments, sunscreens, stress response molecules, and act as substrates for downstream reactions (Winkel-Shirley 2001). The network of enzymatic reactions of the anthocyanin pathway gives rise to three principal classes of red, purple, and blue pigments (the monohydroxylated pelargonidin, dihydroxylated cyanidin and trihydroxylated delphinidin, respectively; Fig. 1, S1). Further downstream modifications of these basic building blocks result in the diversity of colorful anthocyanin pigments found across the flowering plants.

**Fig 1.**
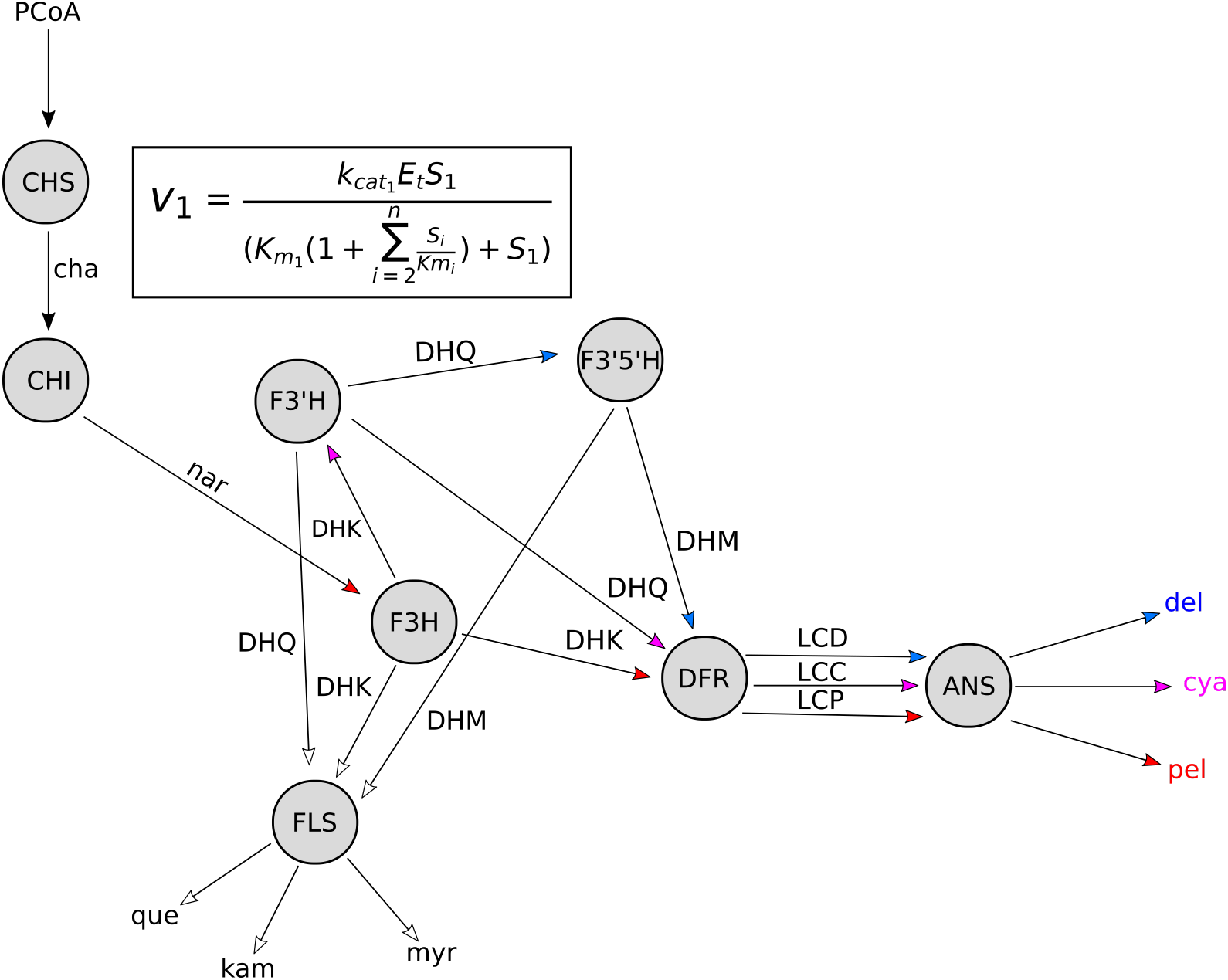
Diagram of the simplified anthocyanin pathway model. The simplified model is depicted as an enzyme-centric pathway diagram, including the key anthocyanin and flavonol branches (see Fig. S1 for classical substrate-centric diagram). Grey circles are enzymes. Arrows indicate influx and outflux of floating species. Arrow placement indicates the linkage: e.g. DHM to LCD to del. Blue arrows lead to delphinidin, red arrows lead to pelargonidin, purple arrows lead to cyanidin. Inset shows general form of the irreversible rate law (accounting for substrate competition): *S*_1_ is concentration of substrate 1, *K*_*cat*, 1_ is the catalytic constant (turnover rate) for substrate 1, *K*_*M*,1_ is the Michaelis constant for substrate 1, the sum is over all other *K*_*M*_ values and concentrations for competing substrates (Schäuble et al. 2013). A complete set of rate laws for all pathway reactions are available in the supplemental text. Floating species abbreviations are: PCoA (P-Coumaroyl-CoA), cha (chalcone), nar (naringenin), DHK (dihydrokampferol), DHQ (dihydro-quercetin), DHM (dihydromyricetin), que (quercetin), kam (kampferol), myr (myricetin), LCD (leu-copelargonidin), LCC (leucocyanidin), LCD (leucodelphinidin), pel (pelargonidin), cya (cyanidin), del (delphinidin). Del, cya, and pel are the anthocyanidins, which are glycosylated to form the various anthocyanins. Kam, que, and myr are the flavonols. Enzyme abbreviations are: CHS (chalcone synthase), CHI (chalcone isomerase), F3H (flavanone-3-hydroxylase), F3’H (flavonol-3’hydroxylase), F3’5’H (flavonoid-3’5’hydroxylase), DFR (dihyroflavonol-4-reductase), FLS (flavonol synthase), ANS (anthocyanidin synthase). The suite of glycosylating and methylating enyzmes (e.g. UF3GT, RT, OMT, (Winkel-Shirley 2001)) that convert the core three anthocyanidins (shown) into the wide range of anthocyanins are not included, for simplicity.

Building on existing knowledge of this pathway, experimental work spanning several decades has identified genetic hotspots for flower color evolution. For example, shifts among the three types of anthocyanins consistently target two branching enzymes (F3’H and F3’5’H), whose activity determines the relative production of three anthocyanin precursors (DHM, DHQ, DHK, Fig.) (Streisfeld and Rausher 2011; Wessinger and Rausher 2012b). These changes may involve structural mutations that alter enzyme function (Wessinger and Rausher 2014), regulatory changes that shift expression (Des Marais and Rausher 2010), or both (Smith and Rausher 2011). The observation of repeated fixation of mutations at certain loci suggests a level of predictability in flower color evolution that may be explained by the underlying pathway structure. At a broad phylogenetic scale, this genetic architecture may lead to evolutionary trends, such as the rarity of some flower colors (Ng and Smith 2018) and the highly asymmetric transition rates among phenotypes (Tripp and Manos 2008). Nevertheless, testing such a relationship requires a theoretical framework that unites the topology of the pathway with phenotypic evolution via changes in the function or expression of pathway enzymes.

The present study takes the first step toward such a unifying framework by creating a mathematical model of the anthocyanin pathway based on classic enzyme kinetics along with a computational pipeline for simulating evolution under directional selection. We use this framework to address four key evolutionary questions: 1) What is the predicted steady-state activity of the anthocyanin pathway based on its topology? 2) How does the pathway control (i.e., which enzymes determine flux) change during evolution toward a new phenotypic optimum? 3) Are certain loci and types of mutations predictably involved in phenotypic evolution? 4) Do trade-offs arising from pathway structure constrain the evolution of pathway components? We hypothesize that the evolution of a new color phenotype will require shifts in pathway control, accomplished by increasing or decreasing the activity of pathway reactions. Accordingly, mutations at loci with the greatest control are predicted to be the greatest contributors to phenotypic transitions (Rausher 2013; Wright and Rausher 2010). However, because of reticulations in the pathway created by the branching enzymes, we also expect that these transitions may lead to trade-offs within and between branches of the pathway that are under selection. The results of these simulation analyses will not only lay the foundation for continued theoretical advances, but will also generate concrete, quantitative predictions that can be tested in empirical systems.

## Methods

### Development and computational implementation of the mathematical pathway model

We used the generalized rate law formulation of (Chou and Talaly 1977; Schäuble et al. 2013) to specify rate laws for each enzymatic reaction in the pathway model (Supplemental text). This formulation scales the Michaelis constant (*K*_*M*_, which relates to binding affinity) for each enzyme-substrate pair by those of competing enzymes to explicitly incorporate substrate competition. In the absence of substrate competition, the rate law reduces to classic Michaels-Menten kinetics (Chou and Talaly 1977; Schäuble et al. 2013). We chose to use irreversible Michaelis-Menten kinetics, because *in vivo* measurements of flavonoid pathway dynamics have been described well by an irreversible model (Groenenboom et al. 2013), but see (Halbwirth 2010; Halbwirth et al. 2006). Moreover, the irreversible form of the model reduces the required number of parameters two-fold relative to the completely reversible equivalent (Schäuble et al. 2013).

Given that detailed kinetic studies are lacking for most anthocyanin pathway enzymes, we designed a naïve model as a starting state to learn about the relative importance of different model parameters and the interplay between them. We chose values for catalytic constants and Michaelis constants from a meta-analysis of enzyme properties in the BRENDA and KEGG databases (Bar-Even et al. 2011; Kanehisa 2008; Schomburg et al. 2002). The model was initialized at a starting state with all parameters of a certain type set to be equal. Values for *K*_*cat*_ (the catalytic rate constant) were set to 14 *s*^−1^, the median of the distribution calculated in Bar-Even et al. 2011. *K*_*M*_ values were set to 0.013 *mM*, one tenth the median value of the empirical distribution to ensure that starting conditions of the model put the upstream substrate concentration well above the *K*_*M*_ of the first enzyme and all subsequent enzymes. Enzyme concentrations were initialized to 0.001 *mM*, a value within the range of naturally-occurring protein concentrations (Miranda et al. 2008). This model represents a naïve starting state, with no preference for any substrates, allowing the “null” unbiased behavior of the pathway topology to be studied. Boundary conditions of the model were imposed by using an upstream source that flows into the pathway at a constant concentration (0.01 *mM*) and sinks at the end of all branches. The sinks are encoded as simple mass action processes with a single rate constant *k*_*sink*_ (0.0005 *M*^−1^) that is the same for all sinks. This process represents the diffusion of the pathway products away from the volume in which they are synthesized, which is consistent with the physiological transport of anthocyanin pigments to the vacuole (Petrussa et al. 2013; Poustka et al. 2007). We implemented this mathematical model of the anthocyanin pathway in Python 3.6 using the Tellurium library (Choi et al. 2016) and the Antimony (Smith et al. 2009) markdown language (see detailed description in supplemental text).

### Evolutionary simulations between defined phenotypic states

We developed a custom Python library, called *enzo* (https://github.com/lcwheeler/enzo), to conduct stochastic evolutionary simulations of the pathway (see Supplemental text for details). Evolutionary simulations are performed using the core PathwaySet object of *enzo*, which acts as a wrapper for the Tellurium pathway model. Within the simulation framework, we randomly select a single parameter from the model at each step and introduce a mutation with effect size randomly sampled from a gamma distribution. Fixation events are determined based on a formula that weights mutations by fitness effect size (see Supplemental text).

Given the structure of the pathway, mutations during these evolutionary simulations can alter any of the enzyme parameters (*K*_*M*_, *K*_*cat*_, and enzyme concentration *E*_*t*_). While these simulated mutations are not modeled at the level of DNA substitutions, they have a close connection with the kinds of variation found in natural systems. The *K*_*M*_ and *K*_*cat*_ for each enzyme are determined by its biophysical properties, which in turn are tied to its protein sequence (Bar-Even et al. 2011; Kaltenbach and Tokuriki 2014; Khersonsky and Tawfik 2010). By contrast, enzyme concentration is controlled by regulation of gene expression and protein degradation (Cooper 2000), and in the case of the anthocyanin pathway, regulation takes place largely at the level of transcription (Quattrocchio et al. 2006). Thus, changes in *K*_*M*_ and *K*_*cat*_ of anthocyanin pathway enzymes likely occur through mutations in enzyme coding sequences (Johnson et al. 2001; Shimada et al. 2001) while changes in enzyme concentration evolve through mutations in cis-regulatory regions or trans-regulatory loci (Sobel and Streisfeld 2013).

In our simulations, each enzyme has a single *E*_*t*_ parameter (reflecting the shared volume of all pathway components in simulations), but can have multiple *K*_*M*_ and *K*_*cat*_ values, depending on the number of substrates. Changes in these parameters occur without correlated effects on other parameters. For example, DFR (Fig. 1) can experience a mutation that improves its activity on one substrate without changing activity on other substrates, consistent with the different *K*_*M*_ and *K*_*cat*_ values for different substrates that have been measured experimentally (Fischer et al. 2003; Katsu et al. 2017; Miyagawa et al. 2015). While it might be biologically reasonable to assume some cost of improvement on one substrate for activity on other substrates (Kaltenbach and Tokuriki 2014; Khersonsky and Tawfik 2010), we surveyed the kinetic data for all multi-substrate anthocyanin pathway enzymes in the BRENDA database (Schomburg et al. 2002) and were unable to detect any consistent patterns of correlated changes, positive or negative. Nevertheless, our model could be expanded to include correlated changes in future.

A total of 9,985 independent evolutionary simulations were run between the naïve starting state and a new optimum, which we selected to be a blue-flowered state. We set the optimum as 90% of the total steady state concentration comprising delphinidin, with a 10% tolerance. All other concentrations were allowed to drift, subject only to the global constraint placed on total steady state concentration of all species, which was held to within a 10% tolerance of that from the initial starting state. These tolerance values are fiexible arguments to the *evolve* function in *enzo* and can be modified to suit the needs of the user.

### Analysis of the simulated data

We designed a series of analyses to address the four questions outlined in the introduction. First, in order to characterize the intrinsic behavior of the pathway, we used metabolic control analysis (MCA), which was developed specifically for analyzing the control of flux through metabolic pathways (Cornish-Bowden 1995; Heinrich and Schuster 1996; Kacser et al. 1995). The standard approach in MCA is to calculate two properties for each pathway reaction: 1) the flux and/or concentration control coefficients, which are the partial derivatives of each flux/steady state concentration with respect to enzyme activity, and 2) elasticities, which are sensitivity coefficients calculated by taking the partial derivative of each reaction rate with respect to each substrate concentration. Together, these measures allow us to determine how pathway enzymes differ in their control over concentrations of floating species (substrates and products) and how reactions vary in their sensitivity to substrate concentration. We used the MCA tools in Tellurium (Choi et al. 2016) to calculate control coefficients and elasticities for our naïve model to assess how the structure of the pathway relates to the steady-state pigment production. We applied the same tools in addressing our second question regarding the changes in pathway control during evolution to a new optimum. In this case, we repeated the MCA analyses on each evolved model and subtracted starting-point control and elasticity matrices from the median-optimal matrices in order to quantify the shift in control.

We next used the simulation data to identify the loci and the types of changes that contributed to evolution toward the delphinidin optimum. For each of our simulations, we counted the number of fixation events involving each model parameter (e.g., enzyme concentration, *E*_*t*_, for DFR) during the trajectories. Our *enzo* package also logged the selection coefficients for each mutation introduced during the simulations, and these were summarized across all trajectories to estimate the distribution of fitness effects (DFE) for each model parameter.

Finally, we used the simulations to assess potential evolutionary trade-offs arising from the pathway structure. We first calculated mean Spearman correlation coefficients (*ρ*) between all pairs of floating species as well as all pairs of model parameters (*E*_*t*_, *K*_*M*_, *K*_*cat*_ values) across all of the simulations. We also compared the distribution of parameter values after the evolutionary simulation with the starting state in order to visualize these correlated changes in pathway activity.

## Results

### Mathematical model reveals behavior of anthocyanin pathway topology

Our naïve starting model, with all enzymes at equal concentrations and with equal kinetic parameters, yielded stable time-course dynamics (Fig. 2a) and resulted in a steady state with species concentrations reaching into the low *mM* range (Fig. 2b). The steady state is characterized by a bias toward products requiring the fewest biochemical reactions (kaempferol and pelargonidin), resulting from the greater accumulation of these products early in the time-course (Fig. 2a). The colored pigments pelargonidin, cyanidin, and delphinidin comprise 32.7%, 10.9%, and 6.6% of the total steady state concentration, respectively (Fig. 2b). This chemical profile would likely result in red or pink floral coloration (Ng and Smith 2016a; Smith and Rausher 2011).

**Fig 2.**
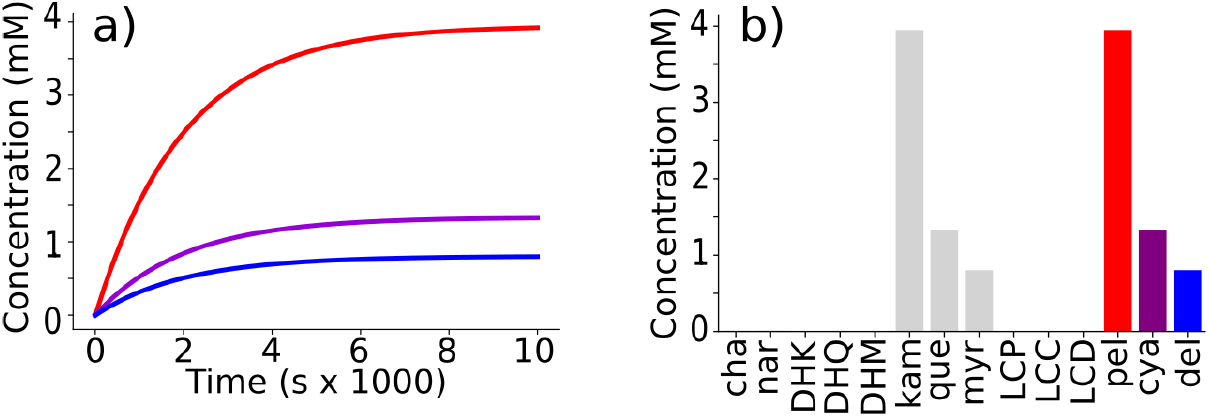
Kinetic behavior of the naïve starting state model. a) time-course simulation of the model showing anthocyanidin concentrations over time: pelargondin (red; highest saturation curve), cyanidin (purple; middle saturation curve), and delphinidin (blue; lowest saturation curve). b) Steady state concentrations of each floating species in the naïve model. The intermediate floating species (e.g. nar and DHM) have a negligible concentration at steady state.

Metabolic control analysis of this naïve model indicated that pathway control is concentrated in the upstream step of the pathway and in the branching enzymes that interact with multiple floating species. The first pathway enzyme, CHS, exerts strong positive control over all downstream reactions and steady state concentrations (Fig. 3a), consistent with previous theoretical studies (Rausher 2013; Wright and Rausher 2010). The control of the branching enzymes varies across products in a manner reflecting their position in the pathway. For example, F3’H has positive control over cyanidin and delphinidin production, which both require its activity, and negative control over pelargonidin, which does not require it (Fig. 3a). Similar patterns of contrasting positive and negative control are seen for other branching enzymes, including F3’5’H, DFR, and FLS. The final enzyme in the pathway (ANS) exhibits little control over any floating species except its substrates, a pattern consistent with its downstream position. The elasticity patterns were similarly intuitive based on pathway structure. For example, CHI and F3H are highly sensitive to the concentrations of their substrates. We also observed negative values due to competition, where, for instance, the reactions involving substrates that compete with DHK (DHQ, DHM, Fig. 1; Fig. S1) are negatively sensitive to DHK concentration (Fig. 3b).

**Fig 3.**
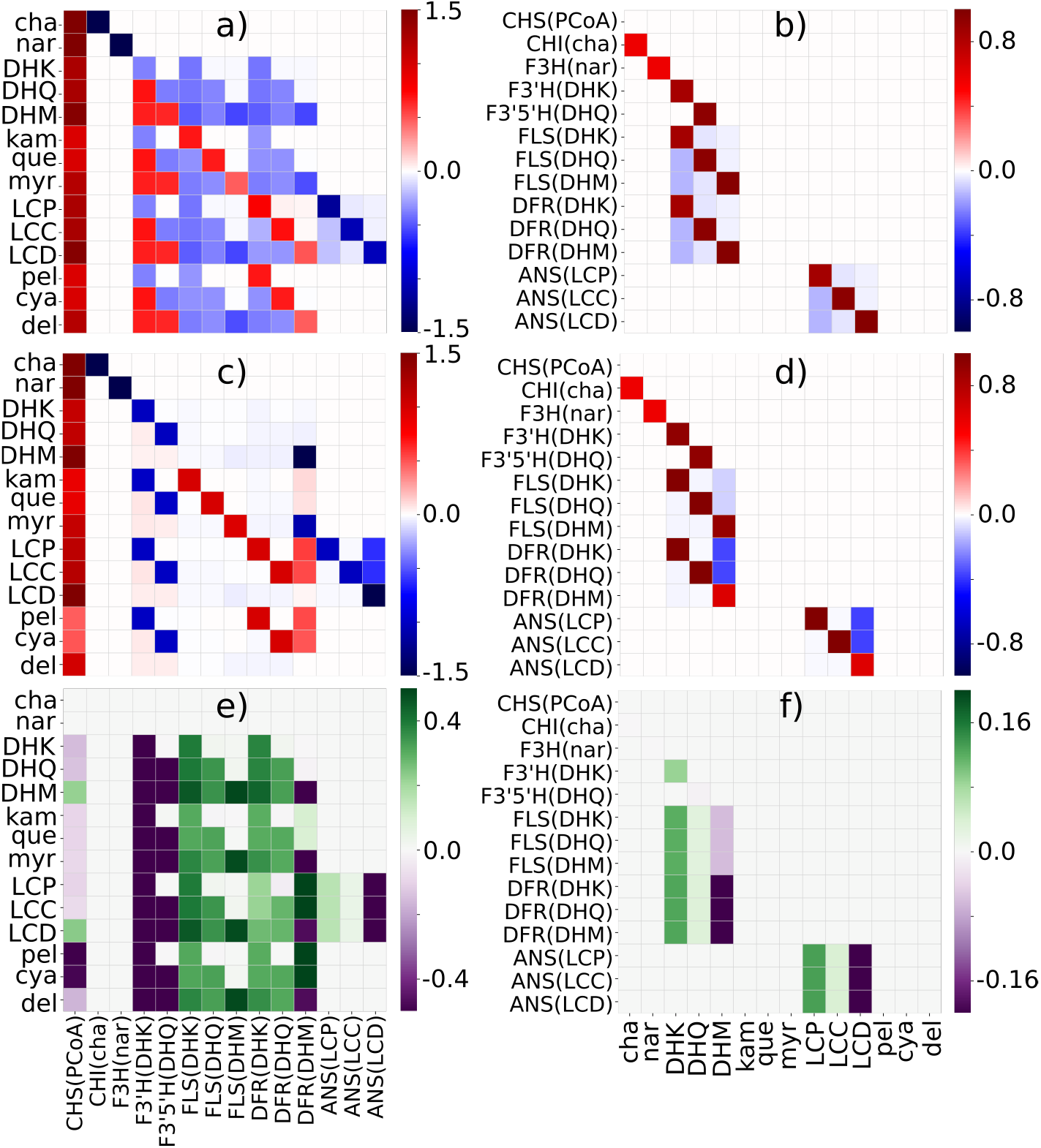
Pathway control shifts to achieve a bias toward the delphinidin branch. a) Heatmap of concentration control coefficients for all pathway reactions (except for “sinks” at boundary conditions) in the naïve starting state model. b) Heatmap of elasticity matrix for all pathway reactions (except for diffusion “sinks” at boundaries) in the naïves starting state model. c) Median optimal concentration control coefficient matrix, taken over all 9,985 simulations. d) Heatmap of median optimal elasticity coefficient matrix, taken over all 9,985 simulations. e) Difference between the two control coefficient matrices (median optimal end-state matrixminus starting-state matrix). f) Difference between the two elasticiity matrices (median optimal end-state matrix minus starting state matrix). In heatmaps, abbreviated floating species identifiers are used for each unique floating species in the model (see Fig. 1). Each unique reaction in the pathway model is denoted using the format “Enzyme(floating species)”, for example ANS(LCD) represents the reaction resulting from ANS acting on LCD as a subtrate. For difference matrices: purple indicates a negative shift in control and green indicates a positive shift.

### Evolution towards delphinidin shifts pathway control

Our evolutionary simulations revealed that the evolutionary transition to delphinidin production required the gain of pathway control by branches leading to or competing with delphinidin. Based on MCA analyses of the 9,985 trajectories that succeeded in reaching 90% delphinidin without error (see Supplemental Results), we observed that F3’H and F3’5’H, both required for delphinidin production, sharply increased in their control coefficients over all downstream substrates and products while the competing enzyme that leads to flavonol production (FLS) lost control over pathway flux (Fig. 3c,e). DFR shifted its activity to gain control over the flux of DHM (the delphinidin precursor), while DFR reactions involving competing substrates (DHQ and DHK) lost control over DHM flux. We found concordant changes in elasticities, with these enzymes becoming more sensitive to the concentration of delphinidin precursors (DHM, LCD) and less sensitive to the competing substrates (Fig. 3d,f). These patterns suggest that while the most upstream gene controls overall flux, the downstream branching enzymes control the type of pigment that is made.

### Loci with greater pathway control are mutational targets

As predicted, we found that approximately 99% of mutations fixed during the simulations involved F3’H, F3’5’H, DFR, or FLS (Table S1), the loci with strong control over delphinidin production. These mutations are spread across all classes of parameters (*K*_*M*_, *K*_*cat*_, and *E*_*t*_), (Fig. S4, Table S1), but for some loci, are concentrated on particular parameters (Table S1 and Supplemental text). Overall, mutations affecting all parameters for the branching enzymes show similar distributions of fitness effects (DFE), from highly negative (*s* = −0.55) to highly positive (*s* = 0.45) (Fig. S3). By contrast, the other pathway enzymes show narrow DFEs skewed towards negative fitness effects (Fig. S3), reflecting that these loci have little control on delphinidin production and that mutations away from the original parameter values are largely detrimental. Across the 9,985 simulations, the median selection coefficient for fixed mutations was 0.044 (range, *s* = 2.4*x*10^−6^ to 0.43), with 90% delphinidin production achieved with a median of 9 mutational steps (*range* = 3 to 80; Fig. 4c).

**Fig 4.**
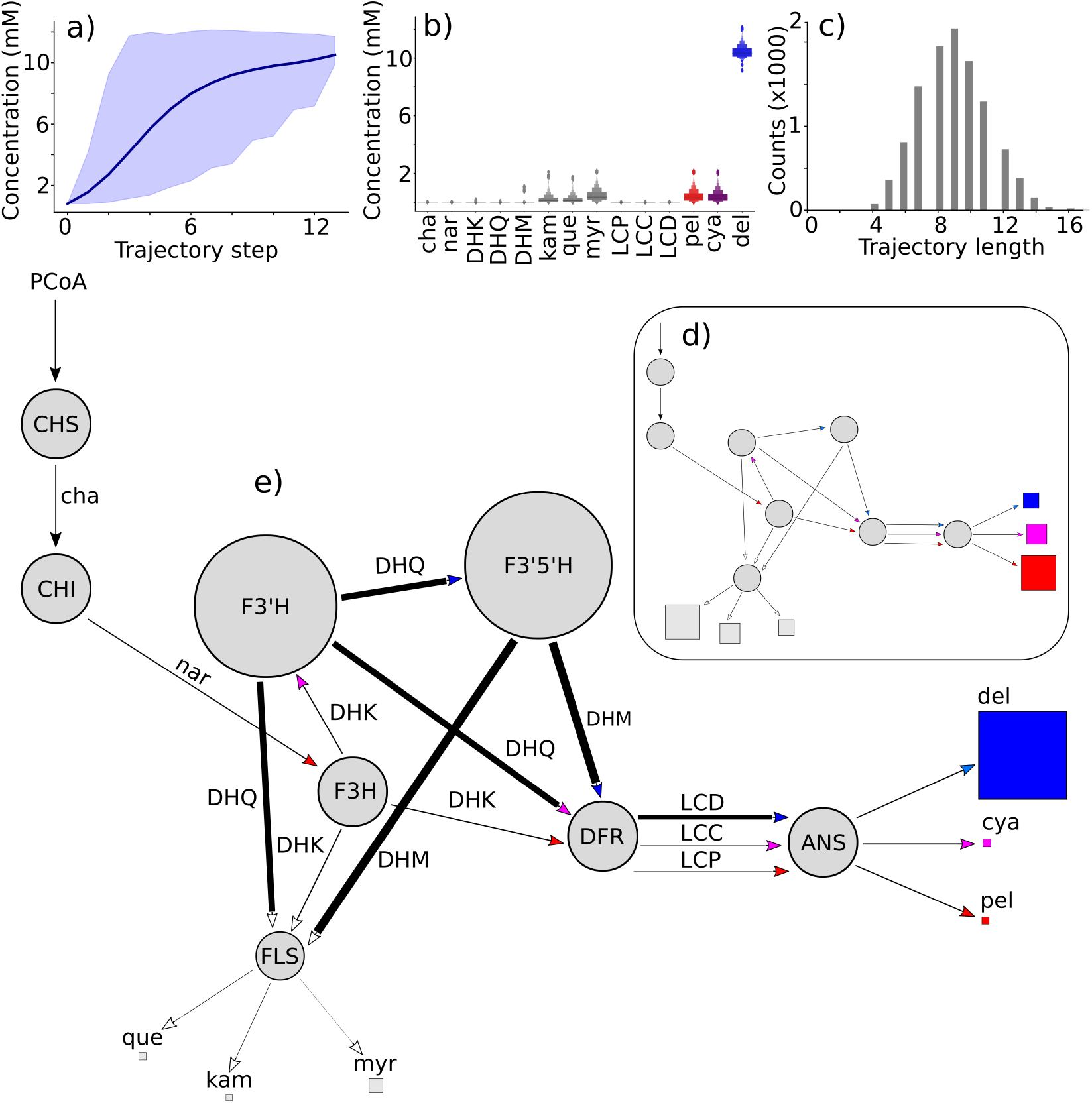
Simulated evolution of delphinidin production. a) Trajectories through delphinidin space. Dark blue line shows mean trajectory, averaged across all 9,985 simulations. Shaded blue area shows envelope containing 95% of trajectories. Top limit is maximum value at each step, bottom limit is minimum value. b) Distributions of optimal (end-point) steady state concentrations for each floating species in the model shown as a boxenplot. c) Distribution of simulated trajectory lengths (median length is 9 steps). d) Inset shows outline of the starting state model, in which all parameters of a given type are equal (see methods). e) The mean values for the model parameters from all delphinidin-optimized end points are shown on the pathway diagram. Enzyme circle size is proportional to concentration. Arrow line thickness is proportional to mean *K*_*cat*_/*K*_*M*_. The area of squares at the end of branches are proportional to steady state concentrations of the indicated floating species produced by a model initialized with mean parameter values.

### Enzyme and substrate competition result in pleiotropic tradeoffs

Evolution toward the delphinidin optimum resulted in sharp and predictable trade-offs in pathway function and activity. Increased delphinidin production came at the cost of decreased production of other pathway products, including the other two anthocyanidins and the three flavonols (Fig. 4d,e, Fig. S2). These effects varied by the number of shared steps, such that compounds requiring the fewest shared steps (namely pelargonidin and kaempferol) showed the strongest decreases. We also observed trade-offs across multiple enzymatic parameters for the loci under selection (F3’H, F3’5’H, DFR, FLS; Fig. S5). For example, the *K*_*cat*_ values for DFR acting on the blue precursor DHM were substantially improved while those for competing substrates (DHQ, DHK) were drastically weakened, resulting in negative correlations (Fig. S5d).

Beyond these antagonistic relationships, we found cases of positive correlations in line with selection toward delphinidin. For example, parameters affecting the same reactions (e.g. *K*_*cat*_ and *K*_*M*_ for DFR acting on DHK) are often positively correlated, whether they are coordinately increasing (because they contribute to delphidinin production) or coordinately decreasing (because they move flux way from delphinidin) (Fig. S5). Similarly, we observed positive correlations between delphinidin production and its two immediate precursors (DHM and LCD) (Fig. S5). Such positive correlations were also seen for other products (pelargonidin, cyanidin) and their precursors (DHK and LCP; DHQ and LCC, respectively). Overall, these patterns reveal how selection on a single pathway product has reverberating effects across all products and precursors due to the changes in amount and activity of their shared enzymes.

## Discussion

In this study, we used a computational approach to probe the fundamental rules governing evolution of floral pigmentation. We constructed a mathematical model of the anthocyanin biosynthesis pathway, which produces a suite of red, purple, and blue pigments. Evolving this model from an unbiased starting state toward a blue-pigmented state revealed the pathway components that are capable of shifting flux between pigment types. These components, in particular the concentration and activity of branching enzymes, were not only the primary mutational targets in our simulations, but also constitute the evolutionary hotspots for transitions among flower color phenotypes in natural systems. We also observed strong trade-offs inherent to the branched pathway topology, which result in predictable shifts in pathway control that are likely to constrain evolution of floral pigmentation. As discussed below, these results have direct implications for connecting color phenotypes with metabolic pathway function and for understanding the evolution of flower color more broadly.

### Intrinsic bias toward red coloration

Our naïve starting state model uncovered an inherent bias toward the production of red pelargonidin pigments, an interesting finding given that they are not the predominant pigments in natural systems. Since this naïve model was based on equal concentrations and equal kinetic parameters for all enzymes, we infer that the tendency towards pelargonidin production (at nearly two-fold the amount of the other two pigments) relates to the topology of the pathway. Specifically, the production of pelargonidin requires five enzymatic steps while the others, cyanidin and delphinidin, require six and seven steps, respectively. This topological bias results in the early accumulation of pelargonidin in the time-course of the model. Consistent with this notion, a similar pattern is observed in the flavonols, where the mono-hydroxylated counterpart of pelargonidin (kaempferol) is produced in greater quantities than the other two flavonols (Fig. 4d,e).

Although we lack an angiosperm-wide dataset on floral pigments, multiple surveys indicate that pelargonidin pigments are generally rare compared to the other classes of anthocyanins. For example, in their survey of 530 species, Beale et al. (Beale et al. 1941) estimated the proportions of taxa containing pelargonidin, cyanidin, and delphinidin-based floral pigments to be 19%, 40%, and 50%, respectively. In a more extreme example, studies in the tomato family suggest that as few as 0.3% of the roughly 2800 extant species produce pelargonidin (Ng and Smith 2016b; Ng et al. 2018). This pattern is not easily explained by genetic mechanisms, as the loss-of-function mutations which often lead to red flowers (Smith and Rausher 2011; Wessinger and Rausher 2015) would instead favor an accumulation of species producing pelargonidin. A more viable explanation is selection driving species away from the pelargonidin-pigmented state, possibly due to pleiotropic effects on the production of the dihydroxylated flavonol quercetin, which is important for UVB protection (Ryan et al. 1998, 2002). By developing a quantitative model linking pathway function to both classes of flavonoids, this study will provide new avenues for testing such hypotheses in future.

### Hotspots of color evolution

Shifts in anthocyanin production, and in turn, flower color, are commonly observed across angiosperms, making these transitions a valuable system for studying the predictability of evolution at the molecular level (Streisfeld et al. 2011; Wessinger and Rausher 2012b). A growing suite of studies has examined evolutionary transitions between flower colorphenotypes, and these implicate two branching enzymes, F3’H and F3’5’H, as the key players determining the type of anthocyanin produced (Des Marais and Rausher 2010; Smith and Rausher 2011; Streisfeld and Rausher 2009b; Wessinger and Rausher 2015). For example, the downregulation of F3’h (with 30-fold to 1000-fold lower expression) was responsible for a switch from blue to red pigmentation in at least four evolutionary cases (Wessinger and Rausher 2012b, 2014). Indeed, a survey of the literature to-date (Table S3) reveals that all fifteen natural examples of transitions between pigment typesinvolved F3’H, F3’5’H or both, while two of these involved mutations in DFR as well. Likewise fifteen of the seventeen examples of engineered pigmentation shifts involved F3’H and/orF3’5’H (Table S3), making thesethe major genetic targets for artificially modifying flower hue(Tanaka Yoshikazu and Brugliera Filippa 2013).Our evolutionary simulations from a red, mostly pelargonidin starting-state to a blue, mostly delphinidin ending-state recovered a similar targeting of changes to F3’H and F3’5’H (Fig. 4e; Table S1; Fig. S4). Mutations at these loci ranged from those altering kinetic properties (enzyme function) to those altering enzyme concentration (analogous to expression) (Fig. 4e, Fig. S4). We observed a mean 5.3-fold increase in F3’H concentration, which combined with a mean 5.3-fold increase in the *K*_*cat*_ for DHK and a mean 1.04-fold tightening of the *K*_*M*_ for DHK, yielded an overall 29-fold mean increase in F3’H activity (Fig. 4; Fig. S4). Changes in F3’5’H activity followed a similar pattern (Fig. 4; Fig. S4). The action of these genes as hotspots for color evolution in nature and in our simulationsmakes sense because, while the upstream loci control flux across the entire pathway, F3’H and F3’5’H control the relative production of the three pigment precursors (DHK, DHQ, and DHM). The preferential fixation of mutations at F3’H and F3’5’H is also in line with theoretical work showing disproportionate occurrence of large-effect beneficial mutations at enzymes with the most control (Fig. 3, (Rausher 2013)). Overall, the striking similarity between these empirical and theoretical studies points to a predictable relationship between the structure of the pathway and loci that underlie phenotypic evolution (Wessinger and Rausher 2012a).

**Fig 5.**
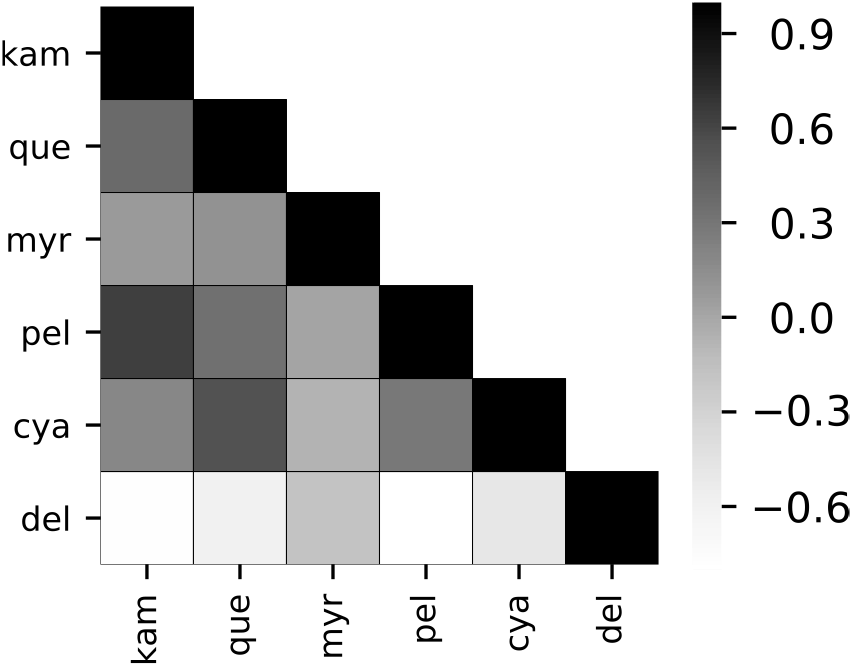
Pleiotropic trade-offs in pathway products. Heatmap of mean Spearman correlations between concentrations of anthocyanins (pel, del, and cya) and flavonols (kam, que, myr) across all 9,985 evolutionary trajectories. Abbreviations are the same as in Fig. 1. A full correlation matrix for all floating species and correlation matrices for model parameters are shown in Fig. S5.

In addition to these major shifts in F3’H and F3’5’H expression and function, we found fixed changes at two other loci, DFR and FLS, which have been implicated less often in natural systems (Table S3). In accordance with selection for increased flux toward delphinidin, DFR improved its activity on the precursor (DHM) roughly 4-fold (on average) during the simulations. Such a matching of flux through the pathway with enzyme activity has been documented in the *Iochroma* system, where DFR from the red species prefers the red precursor DHK while DFR from the blue species prefers DHM (Smith et al. 2013). In this case, the evolution of DFR to track the primary precursor (as determined by F3’H and F3’5’H activity) was interpreted as selection for optimal pathway flux (Smith and Rausher 2011; Smith et al. 2013). Our simulations also recovered fixed mutations that reduced the activity of FLS and directed flux away from flavonol production. Although this gene has been shown to influence floral anthocyanin production in one empirical case (Yuan et al. 2016), its general importance as a locus of color evolution remains poorly understood.

### Evolutionary trade-offs in color production

One of the primary motivations for this study was to understand how the structure of the anthocyanin pathway relates to the phenotypic space for flower color and the mechanisms for moving through that space. Previous surveys have revealed gaps in this color space, i.e. phenotypes that are theoretically possible but not observed in nature (Ng et al. 2018). For example, most taxa produce primarily a single type of anthocyanin, and of those that produce two types, none include the two extremes of the hydroxylation spectrum (pelargonidin plus delphinidin). One explanation for these complex patterns is the presence of trade-offs, which could constrain the course of evolution (Chesmore et al. 2016; Wagner and Lynch 2008). Our study took the first step toward modeling these trade-offs by tracking the changes in pathway function and output during an adaptive walk. These simulations uncovered significant ‘ripple effects’ of selection for delphinidin, which were closely tied to pathway structure. For example, evolution towards the blue state negatively affected production of the other anthocyanins, but the decrease was stronger for the red pelargonidin pigments. This pattern is predicted based on the fact that flux toward delphinidin required higher activity of F3’H and F3’5’H, sapping the DHK precursor for pelargonidin. These types of trade-offs could account for the focusing of production onto one or two anthocyanins as well as the inaccessibility of a phenotype comprising the two extremes (pelargonidin plus delphinidin). Nonetheless, in reaching 90% delphinidin production, we did not observe the complete loss of pelargonidin as is seen in nature (Ng et al. 2018). This discord may reflect several factors that are not incorporated into our simple model, such as the co-regulation of pathway enzymes by shared transcription factors (Albert et al. 2014; Mol et al. 1998).

## Conclusions

This study provides independent support for influence of pathway topology on the distribution of functional mutations for evolving anthocyanin pigmentation. We envision that our model will be valuable for basic research aimed at understanding how selection acts on complex traits as well as applied work focused on flower pigmentation (Sasaki and Nakayama 2015; Shimada et al. 2001) or other phenotypes controlled by metabolic pathways. We note that the basic computational approach developed here could be applied to any enzymatic pathway, opening the door to a broader exploration of how the elements of pathway topology (e.g., length, degree of branching) influence the evolutionary process. It is also straightforward to expand the model to include reversible reactions, multiple ‘containers’ (cellular compartments), pleiotropic mutations, complex selective optima and other features common in biological systems. In the case of the anthocyanin pathway, our ongoing and future research aims to connect this computational model with *in vivo* activity of the pathway and organismal phenotypes, with the ultimate goal of understanding how these molecular mechanisms interact with selection in shaping phenotypic diversity.

## Supporting information

supplemental text updated

## Acknowledgments

We thank members of the Smith lab for useful conversations regarding implementation and interpretation of our computational model. We thank Jacob Stanley and Rutendo Sigauke in the Dowell group at CU Boulder for helpful conversations regarding development of the simulation framework and appropriate implementation of the subsequent analyses. We also thank Boswell Wing, Sebastian Kopf, and the other members of the CU Boulder Geobiology Super Group for their helpful feedback. Finally, we thank Herbet Sauro, Kiri Choi, and Kyle Medley from the University of Washington for their prompt assistance with using the Tellurium library. This work utilized the RMACC Summit supercomputer, which is supported by the National Science Foundation (awards ACI-1532235 and ACI-1532236), the University of Colorado Boulder, and Colorado State University. The Summit supercomputer is a joint effort of the University of Colorado Boulder and Colorado State University.

## Availability of data and materials

The python library *enzo*, is available on github (https://github.com/lcwheeler/enzo). The raw simulated dataset (“wheeler-smith-simulated-dataset.p”) can be found along with other materials on the Zenodo online repository (https://zenodo.org/record/2611739#.XJvOwN-YU8o). Additional details are available in the supplemental text.

## Author contributions

LCW and SDS conceived the study and outlined the computational approach. LCW constructed the mathematical model, wrote the code, performed the simulations, conducted the data analysis, and generated figures. LCW and SDS wrote the manuscript. SDS secured funding for the work. All authors have read and approved the manuscript.

