## supplemental text updated for "Computational modeling of anthocyanin pathway evolution: Biases, hotspots, and trade-offs"

### Computational modeling of anthocyanin pathway evolution: Biases, hotspots, and trade-offs (supplemental text and figures)

Lucas C. Wheeler,[1,*] Stacey D. Smith[1]

1.Department of Ecology and Evolutionary Biology, University of Colorado, Boulder, CO, USA

*

#### Irreversible rate laws incorporating substrate competition

These rates laws encode irreversible Michaelis-Menten kinetics for all enzymes in the pathway. Substrate competition is explicitly incorporated by scaling the $K_M$ of each substrate for multi-substrate enzymes (DFR, FLS, ANS in our model) by the $K_M$ values for all other competing substrates. This rate law form reduces to standard Michaelis-Menten kinetics in the case of a single substrate. In the equations below the parameters are denoted as follows: $K_{cat}$ parameters indicate the catalytic constant, written with the enzyme name in the superscript and the substrate name in the subscript (for example $K^{CHS}_{cat,PCoA}$). $K_M$ parameters indicate the Michaelis constant, written with the substrate name in the superscript and the enzyme name in the subscript (for example $K^{PCoA}_{M,CHS}$). $v$ indicates velocity of the reaction, written with the enzyme name in the superscript and the product name in the subscript (for example $v^{CHS}_{cha}$). Floating species abbreviations are: PCoA (P-Coumaroyl-CoA), cha (chalcone), nar (naringenin), DHK (dihydrokampferol), DHQ (dihydroquercetin), DHM (dihydromyricetin), que (quercetin), kam (kampferol), myr (myricetin), LCD (leucopelargonidin), LCC (leucocyanidin), LCD (leucodelphinidin), pel (pelargonidin), cya (cyanidin), del (delphinidin). Del, cya, and pel are the anthocyanidins, which are glycosylated to form the various anthocyanins. Kam, que, and myr are the flavonols. Enzyme abbreviations are: CHS (chalcone synthase), CHI (chalcone isomerase), F3H (flavanone-3-hydroxylase), F3'H (flavonol-3'hydroxylase), F3'5'H (flavonoid-3'5'hydroxylase), DFR (dihyroflavonol-4-reductase), FLS (flavonol synthase), ANS (anthocyanidin synthase) (Fig.1, Fig. S1).

---

**CHS**

$$v^{CHS}_{cha} = \frac{K^{CHS}_{cat,PCoA} CHS_t PCoA}{K^{PCoA}_{M,CHS} + PCoA}$$

---

**CHI**

$$v^{CHI}_{nar} = \frac{K^{CHI}_{cat,cha} CHS_t cha}{K^{cha}_{M,CHI} + cha}$$

---

**F3H**

$$v^{F3H}_{DHK} = \frac{K^{F3H}_{cat,nar} F3H_t nar}{K^{nar}_{M,F3H} + nar}$$

---

**F3'H**

$$v^{F3'H}_{DHQ} = \frac{K^{F3'H}_{cat,DHK} F3'H_t DHK}{K^{DHK}_{M,F3'H} + DHK}$$

**F3'5'H**

$$v_{DHM}^{F3'5'H} = \frac{K_{cat,DHQ}^{F3'5'H} F3'5'H_t DHQ}{K_{M,F3'5'H}^{DHQ} + DHQ}$$

**FLS**

$$v_{kam}^{FLS} = \frac{K_{cat,DHK}^{FLS} FLS_t DHK}{K_{M,FLS}^{DHK}(1 + \frac{DHQ}{K_{M,FLS}^{DHQ}} + \frac{DHM}{K_{M,FLS}^{DHM}}) + DHK}$$

$$v_{que}^{FLS} = \frac{K_{cat,DHQ}^{FLS} FNS_t DHQ}{K_{M,FLS}^{DHQ}(1 + \frac{DHK}{K_{M,FLS}^{DHK}} + \frac{DHM}{K_{M,FLS}^{DHM}}) + DHQ}$$

$$v_{myr}^{FLS} = \frac{K_{cat,DHM}^{FLS} FNS_t DHM}{K_{M,FLS}^{DHM}(1 + \frac{DHQ}{K_{M,FLS}^{DHQ}} + \frac{DHK}{K_{M,FLS}^{DHK}}) + DHM}$$

**DFR**

$$v_{LCP}^{DFR} = \frac{K_{cat,DHK}^{DFR} DFR_t DHK}{K_{M,DFR}^{DHK}(1 + \frac{DHQ}{K_{M,DFR}^{DHQ}} + \frac{DHM}{K_{M,DFR}^{DHM}}) + DHK}$$

$$v_{LCC}^{DFR} = \frac{K_{cat,DHQ}^{DFR} DFR_t DHQ}{K_{M,DFR}^{DHQ}(1 + \frac{DHK}{K_{M,DFR}^{DHK}} + \frac{DHM}{K_{M,DFR}^{DHM}}) + DHQ}$$

$$v_{LCD}^{DFR} = \frac{K_{cat,DHM}^{DFR} DFR_t DHM}{K_{M,DFR}^{DHM}(1 + \frac{DHQ}{K_{M,DFR}^{DHQ}} + \frac{DHK}{K_{M,DFR}^{DHK}}) + DHM}$$

**ANS**

$$v_{pel}^{ANS} = \frac{K_{cat,LCP}^{ANS} ANS_t LCP}{K_{M,ANS}^{LCP}(1 + \frac{LCC}{K_{M,ANS}^{LCC}} + \frac{LCD}{K_{M,ANS}^{LCD}}) + LCP}$$

$$v_{cya}^{ANS} = \frac{K_{cat,LCC}^{ANS} ANS_t LCC}{K_{M,ANS}^{LCC}(1 + \frac{LCP}{K_{M,ANS}^{LCP}} + \frac{LCD}{K_{M,ANS}^{LCD}}) + LCC}$$

$$v_{del}^{ANS} = \frac{K_{cat,LCD}^{ANS} ANS_t LCD}{K_{M,ANS}^{LCD}(1 + \frac{LCC}{K_{M,ANS}^{LCC}} + \frac{LCP}{K_{M,ANS}^{LCP}}) + LCD}$$

---

### Supplemental methods

#### Anthocyanin pathway model construction and simulation

The simulations were performed under the assumption of a single compartment, analagous to all pathway components being present together in a test tube. We used the default compartment size in Tellurium ($volume = 1$ arbitrary units), which divides out of all the equations used to perform time-course and steady-state calculations. Tellurium numerically integrates the set of coupled differential equations describing the pathway model using the Roadrunner API (Somogyi et al., 2015). To confirm that steady state values calculated by Tellurium were consistent with other standard methods, the model was written out in SBML format ("supplemental-file-1.sbml") (Hucka et al., 2003) and the simulations were then re-run in COPASI (Hoops et al., 2006). The Tellurium and COPASI values for floating species steady state concentrations were identical. All simulations were performed in Python 3.6.3 using the following dependencies: numpy version 1.15.4, tellurium version 2.1.0, pandas version 0.22.0, matplotlib version 2.2.3, seaborn version 0.9.0, and roadrunner version 1.5.1.

#### Evolutionary simulation algorithm implemented in *enzo*

The general outline of the approach taken in this study consists of a few simple steps: 1) write rate laws describing the pathway dynamics, 2) choose a simulation API such as Tellurium (Choi et al., 2016) that is capable of numerically integrating the coupled differential equations to simulate pathway dynamics, and 3) use an evolutionary framework, similar to that which we have implemented, to evolve the pathway model between states.

To perform stochastic evolutionary simulations, we developed a custom Python library, called *enzo* (https://github.com/lcwheeler/enzo). The core Pathway class of *enzo* is a wrapper for Tellurium and Roadrunner model objects, which holds a model object as an attribute. The Pathway object possesses bound methods for conducting evolutionary operations, which allow it to access the built-in attributes of the Tellurium/Roadrunner model. A second PathwaySet class holds an ensemble of Pathway objects, allowing multiple simulations to be conducted while the data for each is collected separately. All Tellurium/Roadrunner model attributes and functions remain accessible to the user. Any Antimony model can be fed to the core PathwaySet object and acted on by the *evolve* method, which evolves a pathway model by selecting on the ratio of one pathway product to the sum of all other products. The number of simulations, replicate Pathway objects held as attributes of the PathwaySet, definition of the optimum state, number of iterations, and other arguments can be provided by the user according to specific needs.

Evolutionary simulations proceed as follows: 1) A Roadrunner model object is initialized from a user-defined Antimony string and held as an attribute of the Pathway object. 2) Steady state concentrations for all floating species in the initial model are calculated. 3) A relative fitness for the current model is calculated compared to the user-defined optimum state. 4) The model enters a loop of user-defined length. 5) For each loop iteration, a single randomly-drawn parameter (from a specified list of available parameters) in the model is randomly mutated by multiplying the current parameter value by a value drawn from a gamma distribution with $\alpha = 0.8$ and $\beta = 3$, selected to make the total number of negative mutations approximately equal to the total number of positive mutations. Any mutation driving total steady state concentration of all floating species outside of a defined tolerance on the starting value (an argument to the *evolve* function) are discarded as non-functional, along with any mutations that result in lack of a steady state solution. 6) Steady state concentrations for the mutant model are computed and used to calculate fitness using the expression: $W = \exp(-(ratio_{current} - ratio_{opt})^2)$, where the ratio is that of the target species steady state concentration to the sum of all steady state concentrations, for the current state ($ratio_{current}$) and the user-defined optimum state ($ratio_{opt}$) (Clark, 1991; Wright and Rausher, 2010; Rausher, 2013). This fitness value is used to calculate a selection coefficient (s) as the difference between the fitness of the current and previous states. 7) For beneficial mutations ($s > 0$), the selection coefficient is used to calculate a fixation probability via an expression $(1 - e^{-s})$, in which population size is ignored for simplicity. This equation weights fixation probability by the mutational effect size. Fixation of the mutation is then determined randomly according the calculated fixation probability. Neutral ($s = 0$) and deleterious ($s < 0$) are discarded, because the fixation probabilities are neglible. This strategy speeds up the simulations by removing an extra step for a large number of mutations, which are more commonly deleterious than advantageous. Each Pathway simulation stops either when it reaches a pre-defined maximum number of mutational steps (50,000) or reaches the new optimum within the user-defined fractional tolerance level for the optimum state.

Steady state concentrations at each fixed step, selection coefficients for each mutation, fixed parameter mutation values, control coefficient and elasticity matrices for each evolved state, and a final optimum parameter set are retained for each simulation and held in memory as attributes of the *enzo* Pathway objects. The entire PathwaySet object, containing the ensemble of simulated models and all of the corresponding data, can be stored as a serialized *pickle* file.

#### Strategy for flower color variation literature review

To compare the predictions of our model against empirical studies of floral color transitions, we performed a thorough review of the available literature. We first used the gephebase.org database (Martin and Orgogozo, 2013) to search for causative mutations underlying changes in floral anthocyanins using the keyword phrases “floral coloration”, “flower color”, and “anthocyanin pigmentation”. We then followed up by manually following citations from the identified articles to find any missed examples. We collected all results for which a transition between floral pigment types had been genetically characterized, counting only entries where the mechanism had been deduced (i.e. the associated changes, regulatory or coding, in anthocyanin pathway genes). We included fixed evolutionary transitions (i.e. at the species or population level), segregating natural variation (i.e. color morphs), and variation in domesticated species. We discarded entries for which the causative mutation was attributed to a transcription factor for which the downstream regulatory effects were unknown, because our model only incorporates information at the level of pathway enzymes. We also discarded transitions where the flower color change involving only differences in the amount of pigment produced (i.e., intensity of pigmentation), as only those which involve changes in pigment type are relevant to our simulations.

We next examined the literature surrounding the metabolic engineering of floral anthocyanin coloration. We used Google Scholar to search the literature for cases where switches between flower colors had been engineered by alteration of pathway enzyme activity. Again, we followed citations from the identified articles to find any literature that was missed in the initial search. We accepted all color-switching transitions involving mutations of anthocyanin pathway enzymes, and again discarded engineered transitions that involved only changes in pigment intensity. The results of this literature search are shown below in Table S3.

### Supplemental results

#### Replicated evolutionary simulations yield an envelope of viable trajectories

We carried out 9,999 replicate evolutionary simulations under the same conditions to yield 9,999 trajectories from the naive starting state to the delphinidin-optimized end-point. Of these, only 9,985 reached the defined optima and did not incur any numerical errors. The failed simulations either 1) reached the maximum number of mutational steps (50,000) without finding the 90% delpinidin optimum within the 10% tolerance; 2) encountered a numerical error in the calculation of the steady state; or 3) encountered a numerical error in the calculation of control coefficient matrices. These errors were detected by the built-in exception-catching scheme of *enzo* and subsequently discarded from the analysis to yield a final set of 9,985 good trajectories.

The median number of fixed mutations to reach the 90% delphinidin optimum was 9 (with a range of 3 to 80 steps). There were 6,077 trajectories with $\leq 9$ steps and only 68 with $\leq 5$ steps (approximately two standard deviations below the mean). All of these very short trajectories were composed entirely of mutations in F3'H, F3'5'H, DFR, and FLS, with a median selection coefficient of 0.14 (and a range of 0.016 to 0.35). 34 of these mutations targeted F3'H, 43 targeted F3'5'H, 56 targeted DFR, and 69 targeted FLS. Overall, there appeared to be a slight bias in the types of parameters targeted by fixed mutations, with 111 in $E_t$, 66 in $K_{cat}$, and 25 in $K_M$, but this pattern seems to vary between enzymes (Table S2).

Collectively, variation in length and mutational composition across simulations resulted in a wide envelope of trajectories when plotted in the anthocyanin space as well as in the spaces of other pathway products (Fig. S2). Because of the 10% tolerance level imposed on the 90% delphindin optimum, there is a small, expected degree of variation around delphinidin concentration at the evolved end-points. The other anthocyanins (pelargonidin and cyanidin) and the flavonol compounds (kampferol, quercetin, and myricetin) have a range of end-point steady state concentrations, from 0% to approximately 10% of total steady state concentration (Fig. 4a).

### Availability of data and materials

The python library *enzo*, written to perform evolutionary simulations of the pathway model, is available on github (https://github.com/lcwheeler/enzo). The scripts used to run the entire simulation procedure ("pathway_sims_final.py" and "run_sims.sh") and a Jupyter notebook containing the subsequent analyses ("complete-analysis.ipynb"), a file containing an SBML formatted version of the starting state model ("supplemental-file-1.sbml"), and the raw simulated dataset ("wheeler-smith-simulated-dataset.p"), in serialized (pickle) format ($> 5GB$) are available in a Zenodo online repository (https://zenodo.org/record/2611739#.XJvOwN-YU8o).

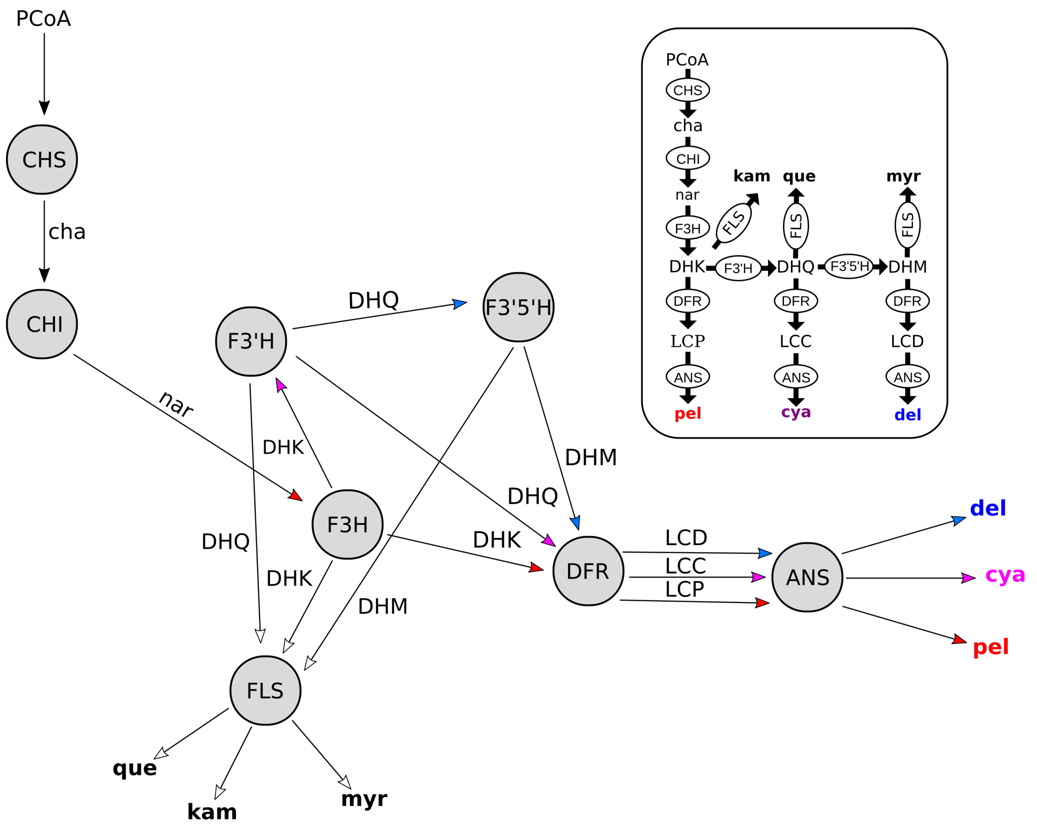


**Fig S1. Alternate diagrams of the anthocyanin pathway.** Enzyme-centric pathway diagram (as depicted in main text) is shown. Inset shows the more typical substrate-centric pathway diagram, with each enzyme depicted multiple times to emphasize the branching structure. Abbreviations for floating species and enzymes follow Fig. 1. Floating species abbreviations are: PCoA (P-Coumaroyl-CoA), cha (chalcone), nar (naringenin), DHK (dihydrokampferol), DHQ (dihydroquercetin), DHM (dihydromyricetin), que (quercetin), kam (kampferol), myr (myricetin), LCD (leucopelargonidin), LCC (leucocyanidin), LCD (leucodelphinidin), pel (pelargonidin), cya (cyanidin), del (delphinidin). Del, cya, and pel are the anthocyanidins, which are glycosylated to form the various anthocyanins. Kam, que, and myr are the flavonols. Enzyme abbreviations are: CHS (chalcone synthase), CHI (chalcone isomerase), F3H (flavanone-3-hydroxylase), F3'H (flavonol-3'hydroxylase), F3'5'H (flavonoid-3'5'hydroxylase), DFR (dihyroflavonol-4-reductase), FLS (flavonol synthase), ANS (anthocyanidin synthase).

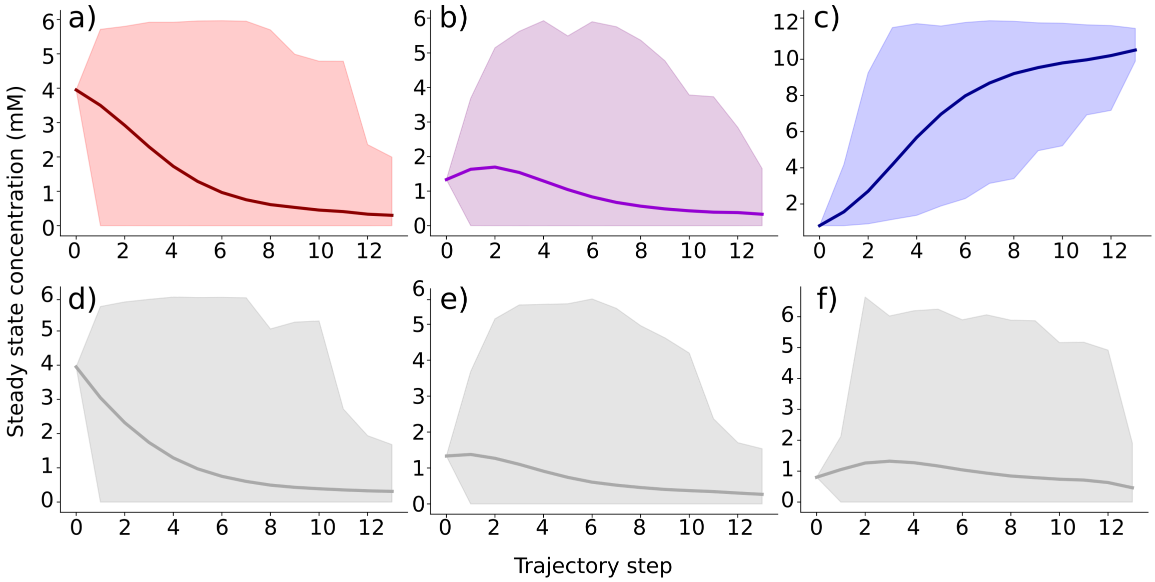


**Fig S2. Trajectories through the six anthocyanin and flavonol dimensions.** Mean trajectories (dark lines) and 95% probability density envelopes (shaded areas) from all 9,985 good simulations are shown for: a) pelargonidin, b) cyanidin, c) delphinidin, d) kampferol, e) quercetin, and f) myricetin.

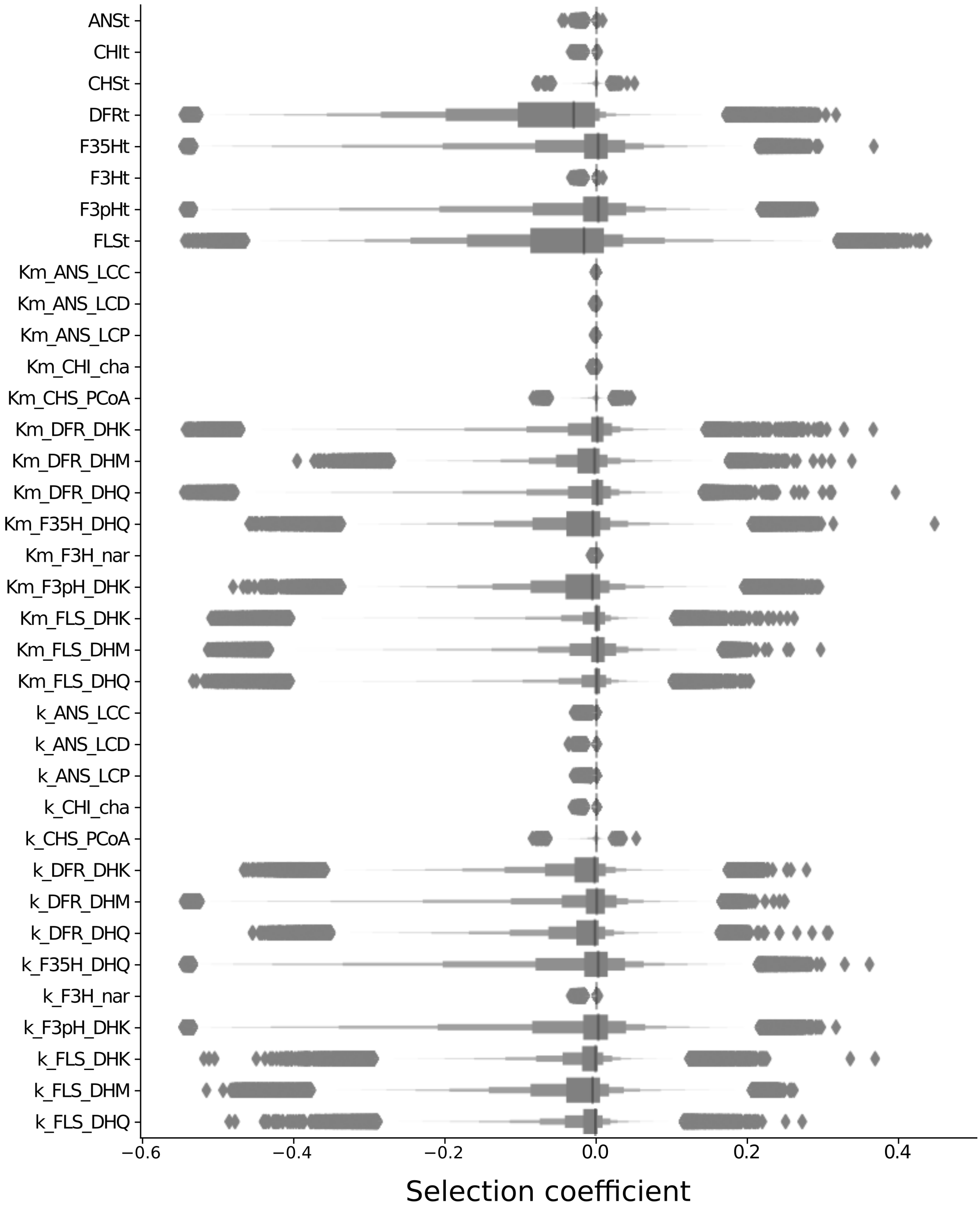


**Fig S3. Distributions of fitness effects vary across pathway enzyme and parameters.** Distributions of fitness effects are shown for each parameter across all 9,985 trajectories, depicted as a boxenplot

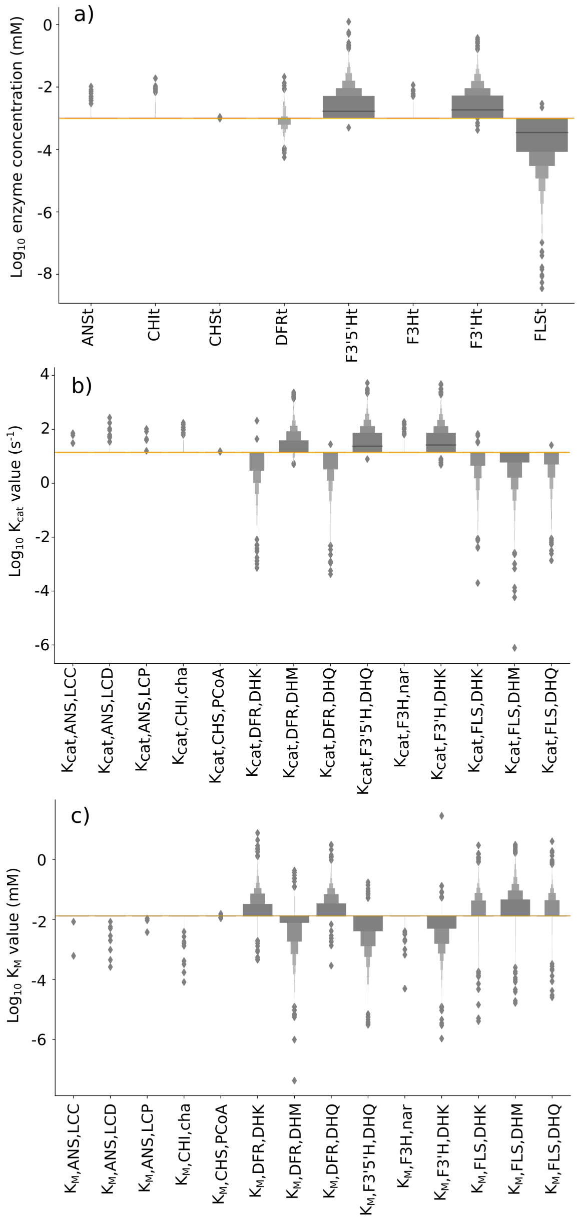


**Fig S4. Parameters of branching enzymes are the dominant components of pathway evolution.** a) Distributions of optimal $E_t$ values from all 9,985 trajectory end-points shown as a boxenplot. Horizontal orange line shows the starting state value. b) Distributions of optimal $K_{cat}$ values from all 9,985 trajectory end-points shown as a boxenplot. Horizontal orange line shows the starting state value. c) Distributions of optimal $K_M$ values from all 9,985 trajectory end-points shown as a boxenplot. Horizontal orange line shows the starting state value.

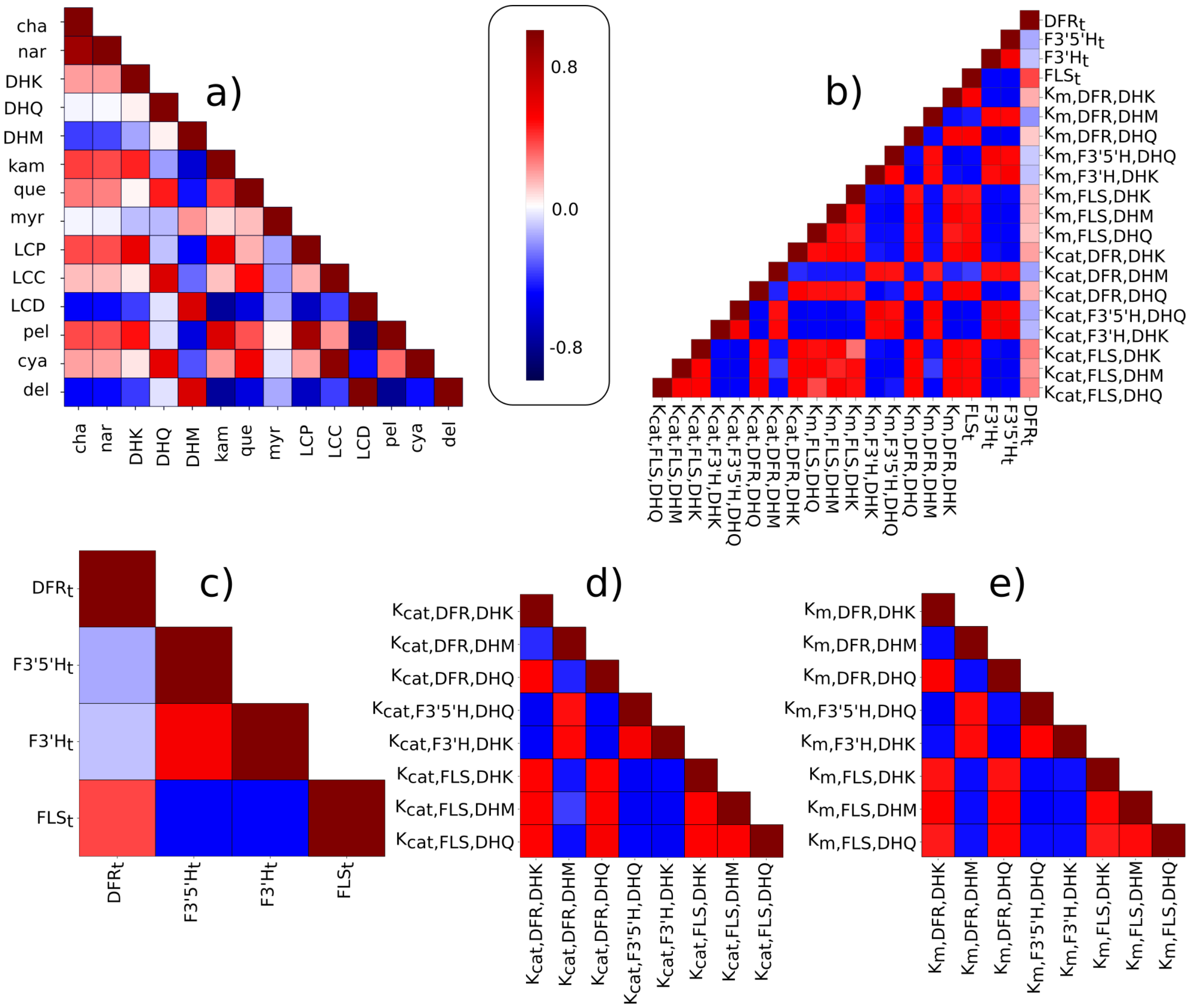


**Fig S5. Pleiotropic trade-offs result from branching and competition.** a) Heatmap of mean Spearman correlations between floating species concentrations across all 9,985 trajectories. b) Heatmap of mean Spearman correlations between parameter changes across all 9,985 trajectories for those parameters accounting for more than 5% of fixation events. Correlations are shown between enzyme concentrations ($E_t$), catalytic constants ($K_{cat}$), and the *inverse* of Michaelis constants ($1/K_M$ in units of $M^{-1}$) to make interpretation more intuitive, because higher $K_M$ (in $mM$ units) corresponds to weaker binding. c) Subset of the parameter Spearman correlation coefficient matrix, showing correlations only between the concentrations of DFR, FLS, F3'5'H, and F3'H. d) Subset of the parameter Spearman correlation coefficient matrix, showing correlations only between the $K_{cat}$ values of DFR, FLS, F3'5'H, and F3'H. e) Subset of the parameter Spearman correlation coefficient matrix, showing correlations only between the $K_M$ values of DFR, FLS, F3'5'H, and F3'H.

| Enzyme (parameters) | Number of fixation events | Percentage of fixation events |
|---|---|---|
| **CHS** | **44** | **0.05** |
| CHSt | 16 | 0.02 |
| Km_CHS_PCoA | 19 | 0.02 |
| K_CHS_PCoA | 9 | 0.01 |
| **CHI** | **100** | **0.12** |
| CHIt | 43 | 0.05 |
| Km_CHI_cha | 19 | 0.02 |
| K_CHI_cha | 38 | 0.05 |
| **F3H** | **83** | **0.10** |
| F3Ht | 37 | 0.05 |
| Km_F3H_nar | 13 | 0.02 |
| K_F3H_nar | 33 | 0.04 |
| **F3'H** | **17560** | **21.8** |
| F3'Ht | 6921 | 8.60 |
| Km_F3'H_DHK | 3714 | 4.62 |
| K_F3'H_DHK | 6925 | 8.61 |
| **F3'5'H** | **17465** | **21.7** |
| F3'5'Ht | 6677 | 8.30 |
| Km_F3'5'H_DHQ | 3988 | 4.96 |
| K_F3'5'H_DHQ | 6800 | 8.45 |
| **FLS** | **24108** | **30.0** |
| FLSt | 7633 | 9.49 |
| Km_FLS_DHK | 2151 | 2.67 |
| Km_FLS_DHQ | 2237 | 2.78 |
| Km_FLS_DHM | 4836 | 6.01 |
| K_FLS_DHQ | 1848 | 2.30 |
| K_FLS_DHM | 3486 | 4.33 |
| **DFR** | **21009** | **26.1** |
| DFRt | 1293 | 1.61 |
| Km_DFR_DHK | 3752 | 4.66 |
| Km_DFR_DHQ | 3725 | 4.63 |
| Km_DFR_DHM | 3112 | 3.87 |
| K_DFR_DHK | 2516 | 3.13 |
| K_DFR_DHQ | 2343 | 2.91 |
| K_DFR_DHM | 4268 | 5.31 |
| **ANS** | **61** | **0.09** |
| ANSt | 20 | 0.03 |
| Km_ANS_LCP | 3 | 0.004 |
| Km_ANS_LCC | 2 | 0.002 |
| Km_ANS_LCD | 8 | 0.01 |
| K_ANS_LCP | 6 | 0.008 |
| k_ANS_LCC | 4 | 0.005 |
| K_ANS_LCD | 18 | 0.02 |

**Table S1. Distribution of fixed mutations for all trajectories.** The observed fixation events across all trajectories decomposed by enzyme and model parameters. The sum of fixations across the parameters of each enzyme are shown in bold. Percentage of total fixation events is listed in the third column.

| Enzyme (parameters) | Number of fixation events | Percentage of fixation events |
|---|---|---|
| **F3'H** | **34** | **16.83** |
| F3'Ht | 15 | 7.43 |
| Km_F3'H_DHK | 6 | 2.97 |
| K_F3'H_DHK | 13 | 6.44 |
| **F3'5'H** | **43** | **21.29** |
| F3'5'Ht | 17 | 8.42 |
| Km_F3'5'H_DHQ | 10 | 4.95 |
| K_F3'5'H_DHQ | 16 | 7.92 |
| **FLS** | **69** | **34.16** |
| FLSt | 68 | 33.66 |
| K_FLS_DHM | 1 | 0.50 |
| **DFR** | **56** | **27.72** |
| DFRt | 11 | 5.45 |
| Km_DFR_DHQ | 2 | 0.99 |
| Km_DFR_DHK | 7 | 3.47 |
| K_DFR_DHK | 19 | 9.41 |
| K_DFR_DHM | 1 | 0.50 |
| K_DFR_DHQ | 16 | 7.92 |

**Table S2. Distribution of fixed mutations for trajectories with five or fewer steps.** The observed fixation events in the 68 very short trajectories ($\leq 5$ steps), decomposed by enzyme and model parameters. The sum of fixations across the parameters of each enzyme are shown in bold.

| Citation | Taxon | Color change | Pigment change | Enzyme Changes | Type |
|---|---|---|---|---|---|
| (Hoshino et al., 2003) | *Ipomea nil, I. tricolor,I. purpurea* | blue/red polymorphism | cya to pel | F3'H (coding mutations) | N |
| (Zufall and Rausher, 2004) | *Ipomea quamoclit* | blue to red | cya to pel | F3'H (downregulation), DFR (specificity) | N |
| (Streisfeld and Rausher, 2009) | *Ipomea udeana, I. quamoclit, I. horsfalliae* | blue to red | cya to pel | F3'H (downregulation) | N |
| (Des Marais and Rausher, 2010) | *Ipomea purpurea, I.pomea quamoclit, I. coccinea, I. ternifolia* | blue to red/pink | cya to pel | F3'H (downregulation) | N |
| (Hopkins and Rausher, 2011) | *Phlox drummondii* | blue to red | del to cya | F3'5'H (downregulation) | N |
| (Mizuta et al., 2010) | *'Oomurasaki' azalea* | purple/red polymorphism | del to cya | F3'5'H (downregulation) | N |
| (Ishiguro et al., 2012) | *Antirrhinum kelloggii* | blue to red | del to cya | F3'H (downregulation) | N |
| (Nakatsuka et al., 2006) | *Gentiana scabra* | blue to pink | del to cya | F3'5'H (deactivation) | N |
| (Matsubara et al., 2005) | *Petunia hybrida* | blue to red/pink | del to cya | F3'5'H (deactivation) | N |
| (Mizuta et al., 2014) | *Rhododendron kiusianum, Rhododendron kaempferi* | red/purple polymorphism | del to cya | F3'5'H (downregulation) | N |
| (Moreau et al., 2012) | *Pisum sativum* | purple to pink | del to cya | F3'5'H (deactivation) | N |
| (Wessinger and Rausher, 2014) | *Penstemon barbatus* | blue to red | del to pel | F3'5'H (downregulation) | N |
| (Wessinger and Rausher, 2015) | *Penstemon labrosus, P. subulatus, P. rostrifloris, P. baccharifolius, P. alamosensis, P. miniatus, P. superbus, P. murrayanus, P. centranthifolius, P. utahensis, P. pinifolius, P. eatonii* | blue to red | del to pel | F3'5'H (degeneration/deactivation) | N |
| (Smith and Rausher, 2011) | *Iochroma gesnerioides* | blue to red | del to pel | F3'H (downregulation), F3'5'H (deletion),DFR (specificity) | N |
| (Freyre et al., 2015) | *Ruellia simplex* | pink/purple polymorphism | pel to del | F3'H (upregulation), F3'5'H (upregulation) | N |
| (Sato et al., 2011) | *Saintpaulia* sp. | purple/pink/blue polymorphism | pel/cya/del variation | F3'5'H (regulatory mutations) | E |
| (Nakatsuka et al., 2007) | *Nicotiana tabacum* | pink to red | cya to pel | FLS (downregulation), F3'H (downregulation), DFR (specificity/transgene expression) | E |
| (Meyer et al., 1987) | *Petunia hybrida* | pink to red | cyanidin to pelargonidin | DFR (specificity/transgene expression) | E |
| (Nakamura et al., 2010) | *Torenia hybrida* | blue to pink | del to pel | F3'5'H (downregulation), F3'H (downregulation), DFR (specificity/transgene expression) | E |
| (Seitz et al., 2007) | *Osteospermum hybrida* | blue to red | del to pel | F3'5'H (downregulation) | E |
| (Noda et al., 2013) | *Chrysanthemum morifolium* | red to violet | pel/cya to del | F3'5'H (upregulated transgene) | E |
| (Fukui et al., 2003) | *Dianthus caryophyllus* | pink to blue | pel to del | F3'5'H (upregulated transgene) | E |
| (Nakamura et al., 2015) | *Rosa hybrida* | pink to purple | pel to cya/del | F3'5'H (upregulated transgene) | E |
| (Katsumoto et al., 2007) | *Rosa hybrida* | red to blue | pel/cya to del | F3'5'H (upregulated transgene),DFR (specificity/transgene expression) | E |
| (Nakatsuka et al., 2008) | *Gentiana triflora, Gentiana scabra* | blue to magenta | del to cya | F3'5'H (downregulation) | E |
| (Ueyama et al., 2002) | *Torenia hybrida* | blue to magenta | del to cya | F3'H (downregulation), FNS (downregulation) | E |
| (Tsuda et al., 2004) | *Petunia hybrida* | purple to pink | del to cya | F3'H (downregulation) | E |
| (Boase et al., 2010) | *Cyclamen persicum* | purple to red/pink | del to cya | F3'5'H (downregulation) | E |
| (Nielsen et al., 2002) | *Eustoma grandiflorum* | blue to purple | del to cya | FLS (upregulated transgene) | E |
| (Shimada et al., 2001) | *Petunia hybrida* | pink to magenta | cya to del | F3'5'H (upregulated transgene) | E |
| (Okinaka et al., 2003) | *Nicotiana tabacum* | pink to purple | cya to del | F3'5'H (upregulated transgene) | E |
| (Brugliera et al., 2013) | *Chrysanthemum× morifolium Ramat.* | pink to purple-blue | cya to del | F3'5'H (upregulated transgene) | E |

**Table S3. Empirical studies of the genetic basis for differences in floral anthocyanin type in natural and engineered systesms.** The literature search used to generate this table is described in the supplemental methods. Natural (N) examples encompass changes that have occurred in the wild along with spontaneous changes in cultivated varieties. Engineered (E) changes are those that have been intentially introduced using biotechnology (i.e. transgenics, gene silencing, etc.). Pigments are abbreviated: pelargonidin (pel), cyanidin (cya), and delphinidin (del) as in figure S1.
